# In Sync With Sound: Interpersonal synchronization as an objective measure of listening engagement

**DOI:** 10.1101/2025.02.26.639799

**Authors:** Lotte Lambrechts, Bernd Accou, Jonas Vanthornhout, Bart Boets, Tom Francart

## Abstract

Listening engagement plays a crucial role in effective communication and knowledge transfer, reflecting a state of deep absorption in an auditory stimulus. The available behavioral assessments of listening engagement are limited in capturing its multidimensional nature. Measuring listening engagement objectively using brain and body signals is a compelling alternative. In particular, interpersonal synchronization—observed through synchronized neural and physiological responses among listeners—may offer a promising objective measure. In this study, we examined synchronization of brain activity, heart rate, and electrodermal activity between participants while they listened to engaging and non-engaging stories. Subjective engagement ratings were collected as a ground truth reference. Results showed higher interpersonal synchronization across all modalities when participants listened to engaging stories compared to non-engaging ones, with significant correlations between synchronization measures and ratings of subjective engagement. These results confirm that synchronization may be a reliable, objective marker of listening engagement. We anticipate that these results will provide a powerful framework for future research in relevant domains, including auditory neuroscience, education, and communication science, and enable impactful real-world applications, such as automatically tracking audience engagement in education, entertainment, and broader societal contexts.

## Introduction

Every second of the day, we encounter various sounds, with speech one of the most pervasive. Where some of these seem negligible (train station announcements), carry little meaning (listening to an automated voice message) or don’t capture you’re interest (listening to a boring lecture), others, like a conversation with a dear friend, will encourage you to be fully present and absorb every detail. Despite these vastly different listening experiences, in all cases our brains will process and interpret these speech sounds, tracking their complex acoustic dynamics [1], [2], [3]. Neural tracking of speech has been established as a reliable and valid measure of natural speech processing and perception, allowing for an objective assessment of speech understanding [1], [4]. Research has shown that this neural representation of speech is modulated by auditory attention to speech [5], [6]. Auditory attention has previously been understood as the conscious decision to direct one’s focus to an external or internal presence [7], [8]. While this has been shown to significantly affect speech processing, it does not completely account for the complexities of everyday listening experiences. Research suggests that when you are genuinely interested and motivated to listen to a speech source, you can enter a state of listening engagement in which you fully absorb everything that is being said. Based on previous research, we have defined listening engagement as deep **immersion in an auditory stimulus, where individuals feel connected to the narrative and its characters** [9]. Listening engagement is different from attention, in that it also involves an unconscious and affective response related to the content or the speaker [9], [10]. It may involve narrative presence [9], [11], a state where listeners lose awareness of their surroundings and feel immersed in the narrative world. Listening engagement yields several beneficial outcomes: It is linked to enhanced speech processing and communication, enhanced memory retention [12], [13], higher listening enjoyment [10], [14], and the experience of a state of flow [9], [15]. Consequently, improving listening engagement could positively impact daily life by for instance enhancing education and media entertainment.

Thus far, listening engagement has mostly been evaluated through behavioral self-report rating scales [9], [11], [16], such as the ‘Narrative absorption scale’ and the ‘Narrative engagement scale’. These well-validated measures can reliably capture variability in listening engagement [17]. Nevertheless, behavioral self-report measures hold inherent limitations, as they may disturb the immediate experience and induce biases (e.g., socially desirable responses when students have to rate their engagement with a lecture). Additionally, they provide only limited temporal resolution, failing to capture moment-to-moment fluctuations in engagement. To address these limitations, an objective measure of listening engagement could provide valuable insights by minimizing bias and allowing continuous assessment. With this research, we will explore how brain and body responses may serve as reliable indicators of listening engagement. Given the strong affective response we expect it to elicit, the combination of brain and body responses may better reflect listening engagement than just brain response alone [18], [19].

In the field of socio-affective neurosciences, it has already been shown that when people share an experience, their brain and body responses will synchronize [20], [21], [22], [23], [24]. Recently, researchers have started to explore the possibility of interpreting listening to the same sound, also as a shared experience. Following this, there is some initial evidence that when people consciously process engaging stories, their physiological responses tend to synchronize [25], [26], [27]. Interpersonal synchronization occurs across multiple physiological modalities, including brain activity, heart rate (HR), and electrodermal activity (EDA). However, considerable uncertainty remains regarding the factors that drive interpersonal synchronization. While it has been observed in direct conversational interactions [28], it also occurs when individuals (jointly) passively listen to the same auditory stimulus [25], [26], [27]. Remarkably, interpersonal synchronization has been found even when participants listen separately to the same stimulus at different moments [26], [27], suggesting that synchronization is not uniquely driven by direct social interaction [25], [29], but rather by shared cognitive and emotional engagement with the content. However, of course, social interactions may amplify these effects.

Various mechanisms have been proposed to explain interpersonal synchrony. Bevilacqua et al. [14] and Madsen & Parra [26], [27] suggest that it arises from the conscious and attentive processing of an engaging stimulus, driven by shared neural oscillations and cognitive processes in response to the content. For example, attentive listening to an engaging stimulus modulates brain activity in a way that aligns across individuals, leading to synchronization. Pérez et al. [25] support this view, proposing that HR synchronizes across participants in response to cognitive processes when individuals listen to the same stimulus. However, other studies suggest that HR may also be influenced by emotional responses elicited by the stimulus, suggesting a broader interplay among cognitive and affective factors [30]. Given that bodily responses such as HR and EDA are closely linked to emotions and arousal [18], [31], [32], emotional processing likely plays a role in listening engagement alongside cognitive mechanisms.

To further understand the relationship between listening and interpersonal synchronization, Madsen and Parra [26], [27] showed that synchronization decreased when participants were distracted from listening through a cognitively demanding task (distracted condition). This suggests that synchronization may be influenced by attention, which is known to be a precursor of engagement. Despite these interesting findings, no conclusions regarding the relationship between synchronization and listening engagement can be validated, as they only manipulated attention and not listening engagement. In addition, previous studies have either lacked a behavioral ground truth [17], [26], [27] or used behavioral measures not directly relevant for engagement [33], making it challenging to ascertain whether the auditory stimulation was genuinely engaging.

The present study aims to address these gaps by employing an objective synchronization measure to distinguish between engaging and non-engaging conditions and to examine the relationship between synchronization and listening engagement. To overcome limitations in prior studies, we meticulously selected engaging and dull stimuli for the participant group and incorporated a behavioral measure of engagement. We hypothesize that interpersonal synchronization will be higher when individuals are engaged with the stimulus and that synchronization levels will predict self-rated listening engagement.

## Materials and Methods

### Participants

Participants subscribed to the experiment in dyads. They came to the lab and participated with someone they knew well (e.g., friends, romantic partners, or family members). After evaluating the inclusion criteria, we included all 40 participants. They had a mean age of 21 years (range: 18 to 25 years). Of these, 4 were male. All participants had normal hearing, confirmed by a digit triplet screening test (SRT = -10 dB SNR) [34] and/ or a tonal audiogram (thresholds < 25 dB HL at all audiometric frequencies ), with a mean Fletcher index of 13 ± 6 dB HL. Other exclusion criteria were attention or concentration problems, neuropathological disorders, communication disorders and language disorders. These criteria were evaluated at the beginning of each session through a screening questionnaire. The Medical Ethics Committee UZ Leuven/Research (KU Leuven) approved the study with reference S68492 and each participant provided informed consent before the start of the session.

### Setup

Testing took place in dyads in a soundproof room (Faraday cage). Participants sat next to each other, approximately 1.35 meters apart, facing each other at a 90° angle (chairs turned 45° towards the loudspeaker). A loudspeaker and monitor were positioned 1.5 meters in front of the participants. The setup was calibrated, and stimuli were presented at 60 dB A, equivalent to an everyday conversation level. See *figure 1* for a full overview.

**Figure 1.**
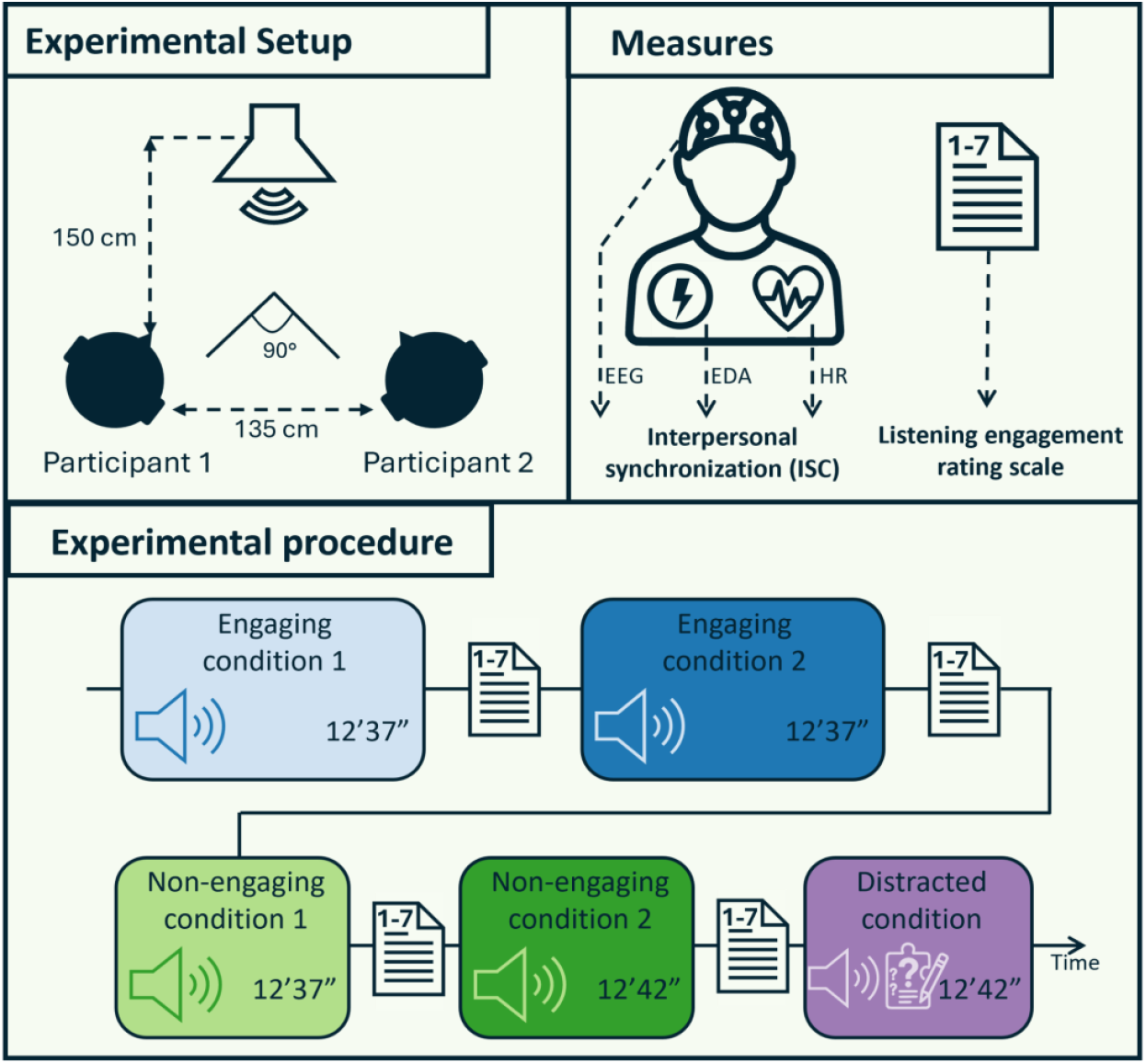
Overview of the experimental setup, the implemented physiological and behavioral measures, and the full experimental procedure. The conditions were presented in a random order but were always administered in their entirety (Engaging – Non-Engaging – Distracted).

### Experimental Design

This study measured brain and physiological responses, as well as subjective listening engagement, in participants exposed to various auditory stimuli.

The experiment consisted of **three different experimental conditions**, each involving the presentation of an auditory stimulus:

#### 1. Engaging condition

Participants listened attentively to a highly engaging stimulus.

#### 2. Non-engaging condition

Participants listened attentively to a stimulus designed to be less engaging.

#### 3. Distracted condition

Participants completed cognitively demanding riddles while an auditory stimulus played in the background, discouraging attention and engagement.

Participants were instructed to listen attentively to the speech stimulus in the engaging and non-engaging conditions. Engagement levels were manipulated by presenting stimuli with varying degrees of intrinsic interest. The **engaging stimulus** was a 46:29-minute episode of a popular Flemish podcast. This stimulus was expected to be inherently interesting, easy to follow, and able to capture individuals’ attention (‘De Volksjury’ episode 19 ‘Maddie McCann’, ranked second place on “Spotify Wrapped” of 2024). The **non-engaging stimulus** comprised excerpts from Dutch debates in the European parliament (25:19 minutes duration), which were self-recorded by one of the researchers. This content was chosen for its neutral/dull and less captivating nature. The engaging and non-engaging stimuli were split into two parts, so we had two engaging and two non-engaging conditions. To minimize the impact of differences between the stimuli on the subject’s responses, we selected stimuli featuring only female voices. After both the engaging and non-engaging conditions, we assessed self-perceived listening engagement, as will be discussed under ‘Behavioral measure - p. 9’. For the **distracted condition**, the participants listened to another non-engaging stimulus (a 12:42-minute self-recorded excerpt of the European Parliament) while solving riddles. Participants were given a new riddle every 2 minutes and were instructed to remember their solutions. After this condition, the solutions were orally discussed. This condition was designed to minimize not only engagement but also attention, serving as a control to differentiate synchronization as a measure of engagement from mere attentional effects.

The full experimental procedure and order of the protocol can be found in *Figure 1*. Each dyad followed the same protocol and same stimuli, with the order of conditions (engaging, non-engaging, and distracted) randomized across dyads to mitigate potential training or fatigue effects.

### Brain and physiological measures

We recorded three objective measures: brain activity via electroencephalography (EEG), EDA, and HR via electrocardiography (ECG). Data were collected using a BioSemi ActiveTwo system (BioSemi - The Netherlands). For EEG, a 64-electrode headcap was applied according to the 10-20 system. We aimed to keep electrode DC offsets (relative to CMS) between -20 and 20 mV. For HR, ECG was measured using two flat electrodes applied at the left lumbar region and below the left clavicle, following established methods [26], [27], [35]. EDA was measured using 2 passive Nihon Kohden electrodes, applied on the pointer and middle finger of the left hand. All signals were recorded at a sampling rate of 2048 Hz, and the recording systems of both participants were synchronized.

### Signal processing

All signal processing (both preprocessing and signal processing) was done in Python (version 3.10), using Numpy [36], Scipy [37] and Pandas (DOI 10.5281/zenodo.3509134).

#### Interpersonal synchronization

Preprocessing was different for each included sensor. All signals were trimmed to match the shortest stimulus length (12:37 minutes) for fair comparison.

##### EEG

EEG was first high-pass filtered using a 4th-order Butterworth filter with a 0.5 Hz cutoff to remove sensor drift. Artifacts exceeding 500 μV were removed, and linear interpolation was performed across the affected intervals. Eyeblink artifacts were removed using a multichannel Wiener Filter [38]. All channels were re-referenced to a common average reference. Lastly, the signal was downsampled to 64 Hz. All filtering steps were done using zero-phase filtering. Additionally, to find the relevant shared information within the EEG from different subjects (for the synchronization analysis), we applied correlated component analysis (corrCA) [39]. CorrCA optimizes weights of a linear combination of EEG channels to maximize correlation between subjects’ EEG and transforms the information to components that are ordered from highest to lowest correlation. These resulting EEG components, calculated from the original 64-channel EEG data are used as input to perform the ISC calculation. To do this, we selected only the first component, as these will be maximally correlated.

##### ECG

From the raw ECG signal, we extracted instantaneous HR. The two ECG leads were re-referenced to each other (ECG1-ECG2) to suppress common-mode noise and enhance the signal-to-noise ratio. We applied the algorithm by Pan & Tompkins et al. [40] to find the R-peaks in the ECG signal. Instantaneous HR was determined as the inverse of the time intervals and interpolated to obtain a sampling rate of 64Hz, similar to the method of Pérez et al., [25].

##### EDA

For EDA, we smoothed the raw signal using a 1.5-second Bartlett window. We computed the difference between consecutive samples to remove low-frequency trends [41], downsampled the signal to 64 Hz and applied a sigmoid transformation (i.e., 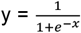, with x = EDA amplitude) to compress large peaks. This scaled the data between 0 and 1, where 0 indicates no change or a decrease, and values above 0 indicate an increase.

To assess interpersonal synchronization, we computed ISCs, which quantify the co-fluctuation of each body response (EEG, HR, EDA) among multiple participants. This involved calculating the average of all the correlations of one participant’s responses with those of all other participants. This resulted in a single averaged correlation value for each participant for each modality for each condition, which could be interpreted as the listening engagement performance of that specific participant in relation to the whole group. A high synchronization value would indicate a high degree of listening engagement, whereas a low value would indicate that the participant was not very involved with the auditory stimulus material. To assess significance, we applied circular shifting for each correlation that was calculated (before averaging). We shifted one of the two signals and then recalculated the correlation. Each value was then Fisher transformed. This procedure was repeated a 1000 times to generate a null distribution, from which we derived a p-value for each synchronization value.

#### Neural tracking of the auditory stimulus

All stimuli were sampled at 48 kHz and normalized for RMS. We extracted the envelope using a gammatone filterbank (Brian2 and Brian2hears toolkits; https://github.com/brian-team/brian2hears), where the sound wave was filtered in 28 subbands between 500 and 5000 Hz. Next, the absolute value of each signal was raised by a power of 0.6 to extract the slow modulations [42]. The envelope was calculated as the average of these 28 subband envelopes. The envelope was downsampled to 64 Hz and used as the stimulus feature in neural tracking. For consistency, all stimuli were trimmed to match the shortest length (12:37 minutes).

No CorrCA was applied for neural tracking analysis. Before analysis, both the EEG data and the speech envelope were bandpass filtered between 0.5 and 4 Hz with a zero phase 4th-order Butterworth filter. To quantify neural tracking, we reconstructed the envelope from EEG data using a linear decoder trained via ridge regression. The decoder had an integration window of 250 ms (or 16 samples), with the regularization parameter (lambda) set to max(abs(EEG covariance matrix)) to prevent overfitting. Linear weights for each channel at different time delays were calculated to minimize the mean squared error between the reconstructed and actual speech envelope. We applied leave-one-fold-out cross-validation, where each recording was split into N non-overlapping folds of 120 seconds. The decoder was trained on all but one fold and then tested on the held-out fold. After training the decoder for a specific stimulus, the envelope was reconstructed from the EEG data and correlated with the speech envelope to obtain a correlation reconstruction accuracy value. This value indicated how well the stimulus was represented in the brain over time.

Statistical significance was calculated for each stimulus separately using circ-shifting *[2]*, which is similar to the procedure described earlier for the synchronization analysis. In this case, the pre-processed speech envelope was randomly shifted in time to generate new noisy correlations. We applied the same Fisher transformation before using all the new correlation values to generate the null distribution.

### Behavioral measure

We evaluated participants’ subjective listening engagement and enjoyment after each engaging and non-engaging active listening condition. We integrated two validated scales, tapping into self-perceived listening engagement. Specifically, we combined the ‘attention’ subscale from the Narrative Absorption Scale [11] and adapted the Engagement Questionnaire by Hannum & Simon [16] so it was more applicable for evaluating listening experiences. This new Listening Engagement Scale consisted of 15 statements about their listening experience, which participants rated on a 1-7 Likert scale. We encouraged participants to be as accurate and honest as possible in their responses. For further analysis, we calculated the mean of these 15 statements for each condition separately per participant. We also administered a short multiple-choice content questionnaire to assess content processing quality and motivate participants to listen actively.

### Statistical analysis

All data visualization and analysis were conducted in R Statistical Software (version 4.3.1, R Core Team, 2023). Statistical significance of behavioral ratings and synchronization measures across different conditions was evaluated using a linear mixed-effects model, with synchronization as the dependent variable, condition as a fixed effect, and participants as a random effect. To determine whether the different conditions had a distinct effect on the synchronization measure, we compared two linear mixed-effects models—one including condition as a predictor and one excluding it—using an ANOVA Likelihood Ratio Test to determine significant differences. We used the following models:

#### Full model

Synchronization value ∼ condition + (1|participants)

#### Reduced model

Synchronization value ∼ (1|participants)

Post-hoc comparisons were corrected for multiple comparisons using a Holm correction. In addition, we calculated Spearman correlations between synchronization values and behavioral ratings. We calculated one correlation per modality across all participants and conditions, each containing 4 data points per participant (two engaging and two non-engaging conditions). Since this violated the assumption of independence, we generated a new null distribution by shuffling the synchronization data by matching the values of each participant to the values of another participant and then again calculating the synchronization-behavior correlation each time. This reshuffling process was repeated a 1000 times. This procedure was applied consistently across all included physiological signals.

To enhance our understanding of the combined physiological responses in relation to the behavioral rating, we computed a summarized index using partial least squares (PLS) regression [43]. Given the high correlation among the different physiological responses, this approach gave us a comprehensive measure while addressing multicollinearity. We applied PLS regression using a leave-one-out cross-validation approach, where the model was trained with three components on data from 39 out of 40 participants, leaving one out for validation. We selected three components as this was the maximum, based on the number of predictor variables [43]. This process was repeated for each participant. We reconstructed the behavioral data using the estimated values from the three PLS components of the regression model. Then we evaluated the alignment between the reconstructed and actual behavioral measures using Spearman’s rank correlation. This final step allowed us to determine how well the PLS regression reconstructed measure aligned with the ground truth behavioral data.

## Results

### Participants rate engaging stimuli higher than non-engaging stimuli on the behavioral Listening Engagement Scale

*Figure 2 (bottom, right panel)* contains the averaged (across all questions) subjective engagement ratings for the different conditions. Due to system failure, we lost responses for the first engaging condition of one participant, the first non-engaging condition of three participants, and the second non-engaging condition of 4 participants. The mean listening engagement score for the engaging condition was 4.8. The mean score for the non-engaging stimuli was 2.1. The linear mixed effects model showed highly significant differences in subjective engagement between both conditions (β = -2.70, SE = 0.098, t = -27.73, p < .001).

**Figure 2.**
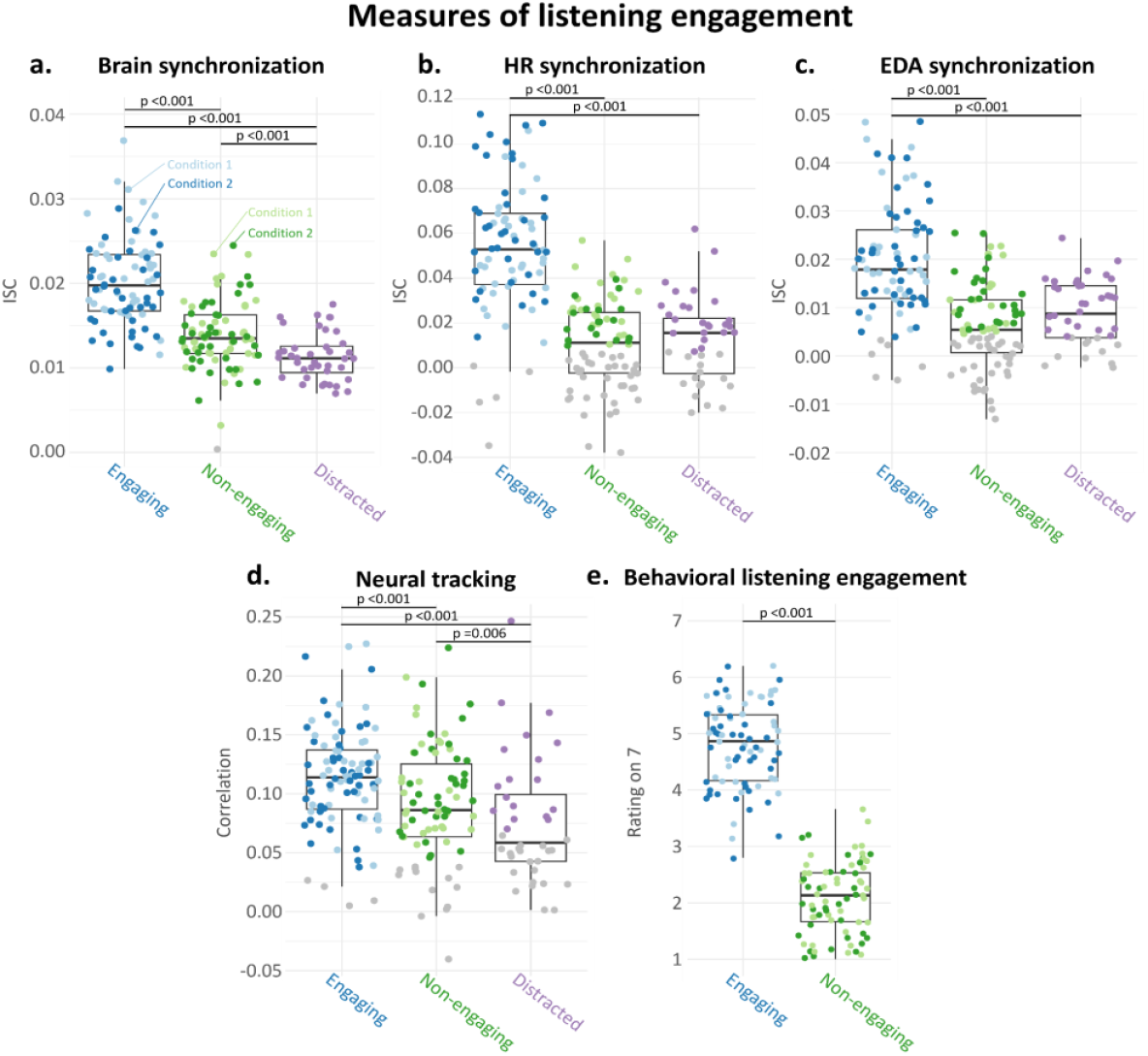
Each data point represents the mean correlation between one participant’s response and all other participants’ responses for a given stimulus and physiological measure. Colored points indicate significance. All measures of listening engagement—both behavioral and physiological (through synchronization)—showed higher engagement in the engaging condition compared to the non-engaging condition. Engaging stimuli were rated significantly higher than non-engaging ones (p < 0.0001). In the physiological data, higher levels of synchronization and neural tracking were observed in the engaging condition relative to the non-engaging condition. Notably, brain synchronization and 119.44, p < .0001), EDA synchronization (χ^2^(2) = 73.392, p < .0001). Post-hoc comparisons with Holm correction (**Table 1**) revealed significantly higher synchronization in the engaging condition (E) than in the non-engaging (N) and distracted condition (D) for all physiological responses. Additionally, brain synchronization was higher in the non-engaging condition than in the distracted condition, but this effect was not observed for HR and EDA. neural tracking also showed significant differences between the non-engaging and distracted conditions.

### Participants display higher interpersonal synchronization when listening to an engaging stimulus

The top three panels in *Figure 2* show synchronization of physiological responses between individuals while listening to auditory stimuli with varying engagement levels. Statistical significance of each data point was calculated using circular shifting.

**Table 1:** Contrasting condition effects for each of the neural and physiological measures.

| | Contrast | $\beta$ (estimate) | SE | t | Adjusted p-value |
| --- | --- | --- | --- | --- | --- |
| Brain | **E - N** | 0.00644 | 0.0005 | 12.878 | **<0.0001** |
|  | **E - D** | -0.00883 | 0.000608 | -14.529 | **<0.0001** |
|  | **N - D** | -0.00239 | 0.000611 | -3.913 | **0.0001** |
| Heart rate | **E - N** | 0.04332 | 0.00356 | 12.174 | **<0.0001** |
|  | **E - D** | -0.04136 | 0.00433 | -9.557 | **<0.0001** |
|  | N - D | 0.00196 | 0.00435 | 0.451 | 0.6525 |
| Electrodermal activity | **E - N** | 0.0145 | 0.00156 | 9.261 | **<0.0001** |
|  | **E - D** | -0.0115 | 0.00190 | -6.036 | **<0.0001** |
|  | N - D | 0.0030 | 0.00191 | 1.571 | 0.1181 |
| Neural tracking | **E - N** | 0.0228 | 0.00471 | 4.839 | **<0.0001** |
|  | **E - D** | -0.0388 | 0.00574 | -6.766 | **<0.0001** |
|  | **N - D** | 0.0228 | 0.00471 | 4.839 | **<0.0001** |

Note. Holm-corrected post hoc comparisons; bold indicates significant contrasts.(E = Engaging condition, N = non-engaging condition, D = Distracted condition)

A highly significant effect of condition could be identified for all physiological responses, based on ANOVA comparisons with and without condition: brain synchronization (χ^2^(2) = 158.24, p < .0001), HR synchronization (χ^2^(2) = 119.44, p < .0001), EDA synchronization (χ^2^(2) = 73.392, p < .0001). Post-hoc comparisons with Holm correction (Table 1) revealed significantly higher synchronization in the engaging condition (E) than in the non engaging (N) and distracted condition (D) for all physiological responses. Additionally, brain synchronization was higher in the non engaging condition than in the distracted condition, but this effect was not observed forHR and EDA.

We also looked at neural tracking results. Significant neural tracking could be found for each condition, but not for every participant, especially in the distracted condition. Using a similar analysis approach as for the synchronization measures, we identified a significant effect of condition for neural tracking (χ^2^(2) = 45.033, p < .0001). Post-hoc comparisons showed significant neural tracking differences between the three conditions (**Table 1**).

### Relation between physiological synchrony measures and behavioral engagement ratings

We analyzed the relationship between the physiological and behavioral data. *Figure 3* presents scatter plots depicting the association between the different interpersonal synchronization values and the scores on the Listening Engagement Scale. Each correlation plot includes data for all participants and conditions involving attentive listening (excluding the distracted condition, as we did not assess any subjective ratings). The first panel contains a scatter plot for brain synchronization, the second for HR synchronization, and the third panel for EDA synchronization. All correlations were significant, indicating a clear relationship between objective synchronization measures and subjective engagement ratings across all modalities. For the different body responses, the scatter plots indicate that higher engagement ratings correspond to higher physiological synchronization between participants’ physiological responses.

**Figure 3.**
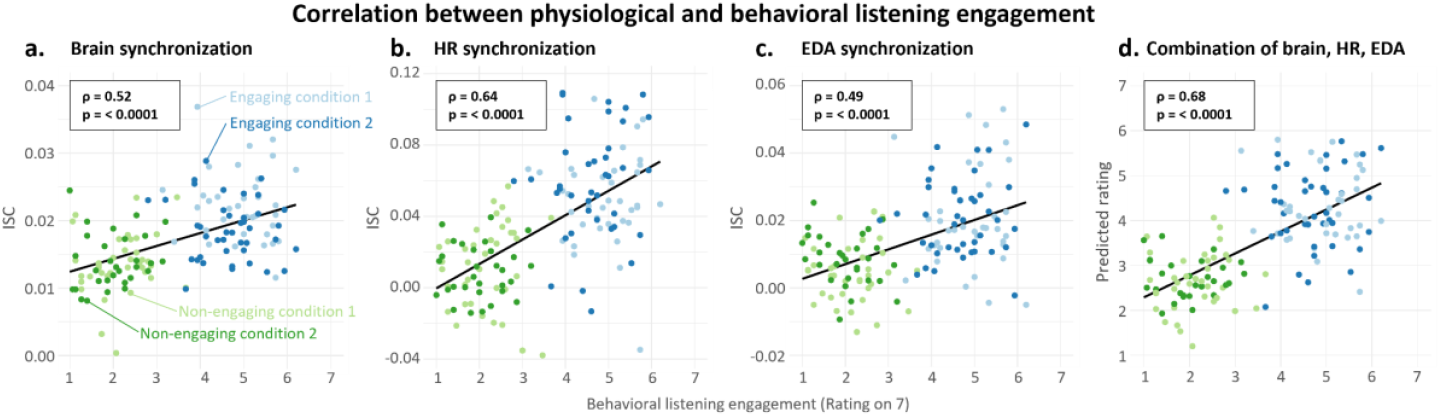
Physiological synchronization (brain, HR and EDA) correlates significantly with behavioral listening engagement ratings. Additionally, a combined physiological measure (predicted through partial least squares regression) also significantly correlated with behavioral listening engagement.

Additionally, we conducted a PLS regression using physiological responses as predictors and behavioral data as the outcome variable. Based on three predictor variables, the regression model explained 88% of the variance in behavioral rating outcomes. To assess how well the behavioral ratings could be reconstructed from these physiological responses, we used Spearman’s correlation to compare the reconstructed data with the Listening Engagement Scale results. This analysis revealed a strong correlation of 0.68 between the PLS measure and the engagement scores, indicating a meaningful relationship between physiological responses and self-reported engagement (see *Figure 3 - last panel*). The observed correlation was higher than any of the individual synchronization statements results; however, no can be made regarding significance.

## Discussion

This paper aims to evaluate interpersonal synchronization as a measure of listening engagement. Twenty duos were invited to our lab and listened to auditory stimuli varying in engagement level (ranging from engaging/interesting to non-engaging/uninteresting). Meanwhile, we recorded physiological brain and body responses. As an extra control condition, participants also completed a distracting task while listening to the stimuli. Our research indicates that physiological synchronization between individuals may be a valuable indicator of listening engagement.

For each of the physiological modalities and across all conditions, significant interpersonal synchronization was observed when people were presented with an auditory stimulus. This finding supports past results that link interpersonal synchronization to conscious speech processing [17], [26], [27]. Extending previous research, our results show that synchronization was higher when participants listened in an engaged manner and significantly lower when the stimulus was non-engaging. This pattern was consistent across all the brain and body signals (EDA and HR), suggesting that engagement activates additional mental processes reflected in physiological responses [44], [45]. These processes might drive interpersonal synchronization across the physiological responses. However, identifying the exact cause of this effect is challenging and requires further research. Nevertheless, our findings expand significantly on previous research, demonstrating that synchrony does not merely result from attentive listening, as participants were encouraged to listen attentively in both the engaging and non-engaging conditions.

We also saw that synchronization declined when participants were distracted compared to when they listened attentively, aligning with previous studies [25], [26], [27]. Through brain responses, including neural tracking and brain-to-brain synchronization, we were able to distinguish between non-engaged listening and distracted listening, indicating that engagement extends beyond attention and conscious processing [17], [26], [27]. These findings support the idea that synchronization can be used as a proxy for listening engagement. However, synchronization did not significantly differ between non-engaged and distracted listening conditions for HR and EDA. From these results, at this point, we cannot make any conclusions as to why this is not the case. It is plausible that attention effects are uniquely cognitively driven and therefore only represented in the brain and that the body responses are more reflective of affective experiences, related to engagement [18], [32]. To the best of our knowledge, this hypothesis has not been tested yet. Further research should be carried out to tease this more apart.

Additionally, we examined the relationship between self-rated listening engagement and objective interpersonal synchronization data. Results revealed strong and consistent correlations between subjective ratings and objective synchronization measures across all three physiological responses. While these correlations may be driven by condition effects, their strength and consistency support the hypothesis that synchronization can predict listening engagement at the individual subject level. PLS analysis revealed a strong relationship between a combined physiological measure and behavioral engagement scores. Although listening engagement is a highly individualized, subjective, and hard-to-capture cognitive-affective construct [9], [10], interpersonal synchronization across these very basic objective physiological measures seems to relate strongly to it. Further analysis could determine whether a single measure, a subset, or the full combination is most effective. At present, the combined measure exhibits the highest correlation, outperforming individual measures. Future research with larger datasets and more advanced models could further refine these findings. Ultimately, these findings could inform the development of sensors and systems capable of automatically monitoring and tracking audience focus and engagement, with promising applications in society, education and entertainment.

While encouraging, these results stem from a deliberately sharp contrast in stimulus types. To maximize the contrast between engaging and non-engaging stimuli, we selected content that was either highly interesting or very dull. However, in everyday life, auditory input often falls along a more nuanced spectrum. Future studies should incorporate a broader range of engagement levels to determine whether a continuous gradient in synchronization can be observed.

Listening engagement is a multidimensional construct influenced by various factors, making it challenging to control all potential confounds [9], [10]. For example, different speakers were used across the conditions, and individual preferences for voice timbre could have affected engagement. To partly control for this, we ensured all speakers were female, eliminating potential effects of the speaker’s sex. However, factors such as fatigue, anxiety, and personal interest [7], [9], [10] may also play a role. Future research should also further investigate how speaker engageability characteristics (e.g., speech rate, prosody, vocal quality) influence listening engagement.

Another key challenge is differentiating between listening engagement and auditory attention. In all active listening conditions, participants were instructed to listen and focus attentively to the presented stimulus. To ensure compliance, participants were informed beforehand that they would be asked to answer multiple-choice questions. While this design theoretically ensures comparable attention levels across both listening activities, engaged and attentive listening may be more intricately linked in practice. As a result, it remains unclear how much of the observed synchronization differences between conditions were driven by engagement or variation in attention. Future studies should further disentangle these overlapping constructs.

Finally, through future analyses, we will examine dyadic effects within these data to gain deeper insights into interpersonal synchronization dynamics and explore whether synchronization is enhanced by physical proximity and real-life presence.

## Conclusion

In this study, participants listened to auditory stimuli while we measured their brain activity, HR, and EDA. The stimuli were designed to be either engaging or non-engaging. Our analysis revealed that interpersonal synchronization was significantly higher in the engaging condition across all physiological responses. Additionally, individual differences in interpersonal synchronization closely correlated with subjective listening engagement scores. These findings strongly suggest that interpersonal synchronization can serve as an objective marker of listening engagement.

